# One Health in Eastern Africa: No barriers for ESBL producing *E. coli* transmission or independent antimicrobial resistance gene flow across ecological compartments

**DOI:** 10.1101/2024.09.18.613694

**Authors:** Patrick Musicha, Mathew A Beale, Derek Cocker, Fiona A Oruru, Allan Zuza, Chifundo Salifu, George Katende, Sylvia Nanono, Fred Isaasi, Kondwani Chidziwisano, Lawrence Mugisha, Henry Kajumbula, David Musoke, Tracy Morse, Shevin Jacob, Nicholas A Feasey, Nicholas R Thomson

**Author notes:** **Corresponding Authors:** Patrick Musicha Nicholas R Thomson.

## Abstract

The One Health paradigm considers the interdependence of human, animal and environmental health. In high-income countries, limited evidence has been found from recent studies to support the importance of a One Health approach to addressing spread of antimicrobial resistance (AMR). Given AMR is a global threat, and we are all interconnected it would be important to know if closer interaction of humans with animals and the environment in low-income present a contrasting picture. We used whole genome sequencing to investigate the genomic diversity and to infer transmission of extended spectrum beta-lactamase producing *Escherichia coli* (ESBL-Ec) between different ecological niches (humans, animals and the environment). We found high diversity of ESBL-Ec with 172 genomic clusters and 167 sequence types identified from 2,344 genomes. Common ESBL genes, *bla*_CTX-M-15_ (67.6%) and *bla*_CTX-M-27_ (14.2%) were carried on a complex network of different plasmids, presenting multiple pathways for dissemination and revealing the high force of selection. Using fine-scale genomic clustering across multiple thresholds ranging from 0 to 20 single nucleotide polymorphisms, we found that genomes isolated from humans, animals and the environment formed overlapping clusters, indicating recent ESBL-Ec transmission and co-circulation both within and between ecological compartments. These findings demonstrate that the One Health approach is highly relevant to tackling AMR in low-income settings, and therefore critical to consider if we are to address the rise of AMR globally.

## Introduction

*Escherichia coli* are found ubiquitously as commensals in the intestinal tracts of humans and warm-blooded animals, but can also cause a range of diseases. In humans, these diseases include both diarrheal and extra-intestinal infections such as urinary tract and bloodstream infections, ^1,2^ which can be life-threatening especially in vulnerable populations including neonates and immune-compromised individuals.^2^ The threat to human health posed by *E. coli* is exacerbated by the spread of antimicrobial resistance (AMR). A recent study on the global burden of AMR estimated 1.3 million human deaths to have been attributed to AMR in 2019, and identified *E. coli* as the leading aetiologic agent of AMR attributable deaths.^3^ Of major concern are extended spectrum beta-lactamase producing *E. coli* (ESBL-Ec), which are resistant to third generation cephalosporins (3GC) and identified by the World Health Organization to be among pathogens of critical priority.^4,5^ In recent years, 3GC resistant (3GC-R) *E. coli* have become more common in all world regions including Africa.^6^ While evidence on burden of 3GC-R *E. coli* infections in Africa remains sparse, there are growing indications they cause higher mortality and morbidity risk than 3GC susceptible *E. coli* infections.^7,8^

Discerning the full diversity of AMR bacteria across different ecological niches and geographical settings is important for identifying reservoirs and sources of AMR, and how AMR bacteria are transmitted between those ecological niches and settings.

Our current understanding of ESBL-Ec diversity is, however, not only limited to studies biased towards samples from human and healthcare settings but also lacks comprehensive geographical and temporal representation.^9^ Recognising that the health of people is closely connected to the health of animals and our shared environment, One Health proposes a holistic approach to designing and implementing interventions for combating health threats. However, while the One Health approach is particularly viewed as important for AMR, the evidence to support it is limited. A study investigating the genomic epidemiology of ESBL-Ec from livestock farms, retail meat and *E. coli* bacteria associated with bloodstream infection in the United Kingdom (UK) found that bacterial lineages and AMR genetic determinants of human clinical isolates were distinct from lineages and AMR determinants of isolates from livestock samples.^10^ Similarly, another study investigating transmission dynamics of carbapenemase-producing *Klebsiella pneumoniae* in Italy reported limited transmission between humans, animals and the environment.^11^ The limited transmission between humans and animals in high income countries was also shown to be true for other Enterobacteriaceae pathogens with different AMR profiles such as *Salmonella enterica*.^12^ Importantly, these studies have posed questions around the degree to which controlling the spread of AMR truly requires a One Health solution, in their settings. However, they also highlighted the pressing need to test the generality of their observations in multiple other environments by conducting research with targeted genotypic data sets across veterinary and public health sectors, especially in different agricultural, political, and sociological conditions.^12^

Here we hypothesised that closer interaction of humans with animals and the environment in settings not protected by adequate environmental sanitation systems, as is often the case in low-income settings, would present a contrasting picture to what has been observed in high-income settings.^13^ Using longitudinally collected household-linked samples from Malawi and Uganda, we investigated the diversity of ESBL-Ec genomes and ESBL determinants and inferred potential transmission events between human, animal and environmental compartments under the Drivers of AMR in Uganda and Malawi (DRUM) study.

## Results

### Genomic diversity and population structure of ESBL-producing *E. coli*

We sequenced 2,828 isolate genomes (2252 [79.6%] from Malawi and 576 [20.4%] from Uganda) identified as likely ESBL-Ec. After performing quality filtering on the raw Illumina reads and genome assemblies, 2,344 (82.9%) of the sequenced genomes were identified as high-quality *E. coli* genomes, including 160 *Shigella spp.*, and all these were included in downstream analyses. Here, we treated the *Shigella* spp. as a specialised *E. coli* pathovar and included them in the analysis due to their phylogenetic indistinguishability from *E. coli.*^14^ There were 1,814 (77.4%) genomes from Malawi, of which 907 (50.0%) were from human stool, 221 (12.2%) from animal stool and 686 (37.8%) from the environment and 530 (22.6%) genomes from Uganda including 380 (71.7%) from human stool, 147 (27.7%) from animal stool and 3 (0.05%) from the environment.

The pan-genome for the 2,344 isolates comprised 23,677 genes, of which 3,066 (12.9%) were identified in at least 99% of the genomes and thus considered to be core. We inferred a phylogeny from 268,606 single nucleotide polymorphism (SNP) sites within our core gene alignment showing the ESBL-Ec from the DRUM study to be highly diverse, with multiple phylogenetic clusters including genomes comprising isolates from both Malawi and Uganda, and from the three ecological compartments we sampled from (Supplementary figure 1). Analysis of population structure using PopPUNK assigned the 2,344 ESBL-Ec isolates to 167 lineages. Of note despite the apparent high diversity in lineages, the distribution of genomes was highly skewed, with approximately 71% (1,675/2,344) of the isolate genomes belonging to only 16/167 (10%) of the ESBL-*Ec* lineages. These sixteen high frequency lineages comprised between 40 and 313 isolate genomes each (Figure 1A). Conversely, most of the lineages (110/167, [65.9%]) were uncommon, containing less than five genomes each.

**Figure 1:**
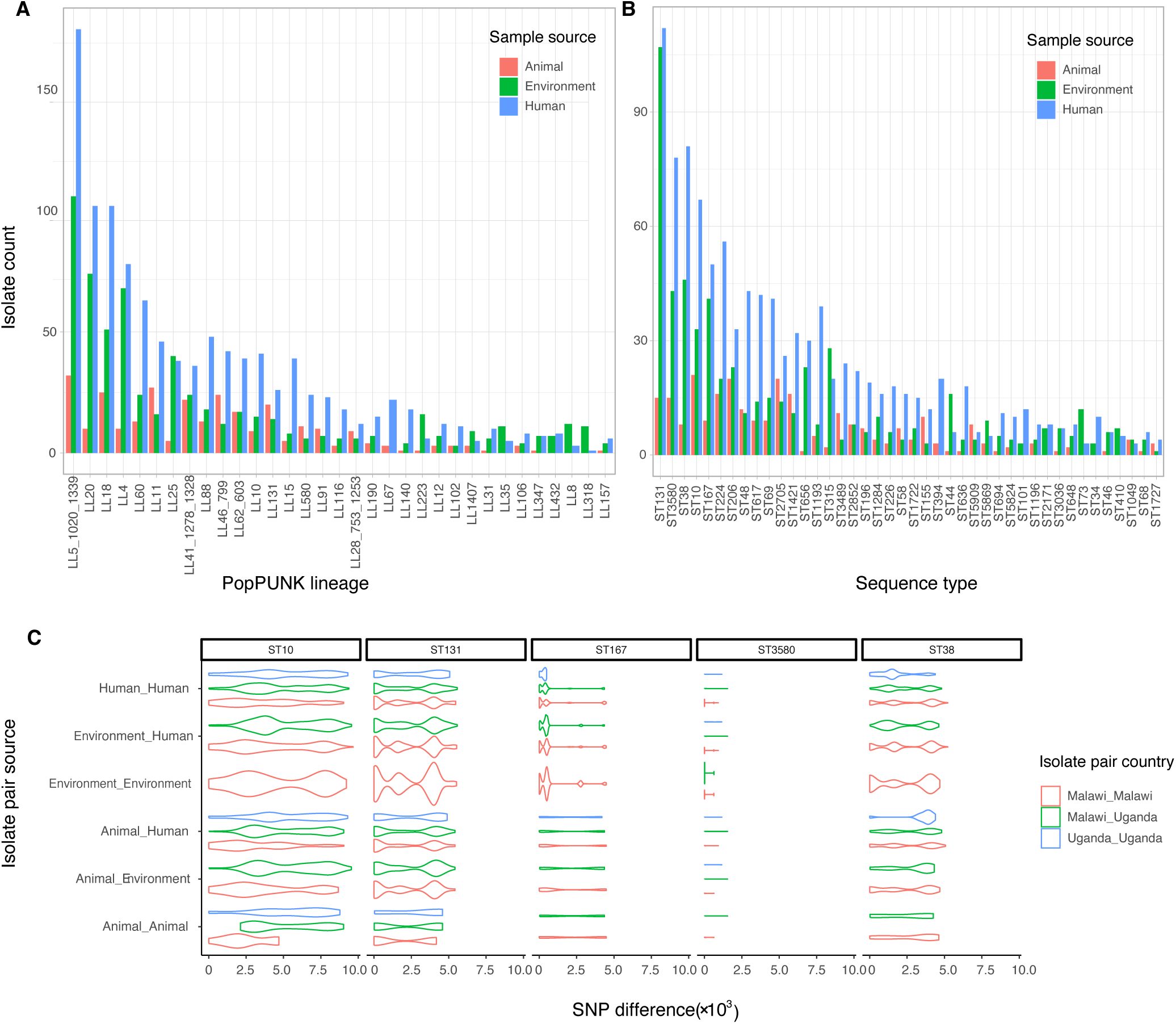
ESBL producing E. coli isolates from Uganda and Malawi are highly diverse. Analysis at the population level (Genomic clusters defined using both PopPUNK and Multi-locus sequence types) indicates a large number of genomic lineages (A) and sequence types (B), in both cases the majority of which are shared by all ecological niches. Panels A and B include PopPUNK lineages and STs with a frequency of at least 10 genomes. Analysis of pairwise SNP distributions within STs (C) shows both close (including identical genomes) and distant genomic relationships. The distribution of pairwise SNP distances within STs by countries of origin and ecological sources of isolate pairs for five most common STs in C, illustrates the high diversity within most ST and the need for high resolution analysis beyond ST level to infer transmission..

To link to the global epidemiology of ESBL-Ec, we typed all isolates using multi-locus sequence typing (MLST) and compared the results to the PopPUNK lineages. As expected, the high diversity was replicated by MLST (Figure 1B), whereby 2239/2344 ESBL-Ec were typed into 170 known sequence types (STs). Fifty percent (85/170) of STs were either singletons or contained just two genomes. Of the STs, ST131 was the most common (n=234/2344, [10.0%]), followed by ST3580 (n=136/2344, [5.8%]), ST38 (n=135/2344, [5.8%]), ST10 (n=121/2344, [5.2%]), and ST167 (n=100/2344, [4.3%]) (Figure 1). Although most common STs and PopPUNK lineages included isolates from both Malawi and Uganda, there were significant differences in ST distribution between the two countries (simulated p-value <0.001, Fisher’s exact test), with ST131 (n=209, [11.52%]), predominating in Malawi, and ST10 (n=45, [8.5%]) predominating in Uganda (Supplementary figure 2). We also found 105/2344 [4.5%] genomes that were not present in existing MLST databases; these genomes were submitted to Enterobase.^15^ Of note, the majority of new STs were related to the ST10 (28/105, ST155 (19/105) and ST131 (7/105) clonal complexes, indicating that they are closer relatives of these dominant STs (Supplementary Table 1).

### Fine-scale lineage within ST diversity and phylogenomic analysis

Within individual STs, we examined the number of core gene SNPs and found this varied, with some STs showing very low diversity (all genomes separated by no more than 5 core SNPs), whilst other STs had very high diversity (some genomes separated by more than 1.0 × 10^!^ core SNPs) (Figure 1C & Supplementary figure 3). Due to these wide variations, STs provided insufficient resolution for investigating the relative diversity and potential transmission of isolates from the different ecological niches and settings (Figure 1 and Supplementary figure). Therefore, to gain better understanding of the population structure and diversity of ESBL-Ec, we combined long (PacBio) and short (Illumina) reads to construct high quality hybrid reference genomes for five STs, present in both countries and including at least 100 genomes (ST10, ST131, ST167, ST38 and ST3580). We mapped short reads for all other genome sequences to call SNPs and accurately determine within ST diversity. Next, hierarchical Bayesian Analysis of Population Structure (hierBAPS) was used to subdivide the STs into hierBAPS lineages (Figure 2): ST10 isolates were subdivided into five hierBAPS level1 lineages. The largest ST10 lineage (L1.GC3) consisted of closely related genomes predominantly from Malawi. The remaining four ST10 clusters consisted of less related genomes, from both Malawi and Uganda, that were characterised by deep-rooted branches (Figure 2A). ST131 and ST167 were each delineated into four hierBAPS level 1 lineages and ST38 into five lineages (Figure 2A-D). ST3580, which was separated into three hierBAPS lineages, was the most clonal (of the top five), with the two most divergent genomes having a genetic distance of 94 SNPs (Figure 2E). Unlike ST10, all the BAPS genomic clusters of ST131, ST167 and ST38 and 3580 comprised of closely related genomes characterised by short branch lengths. Across all the five STs, there was strong phylogenetic mixing of isolates from all the three ecological compartments, but although interspersing of isolates from the two countries could be observed, we also found phylogenetic clustering of genomes by country of origin (Figure 2).

**Figure 2.**
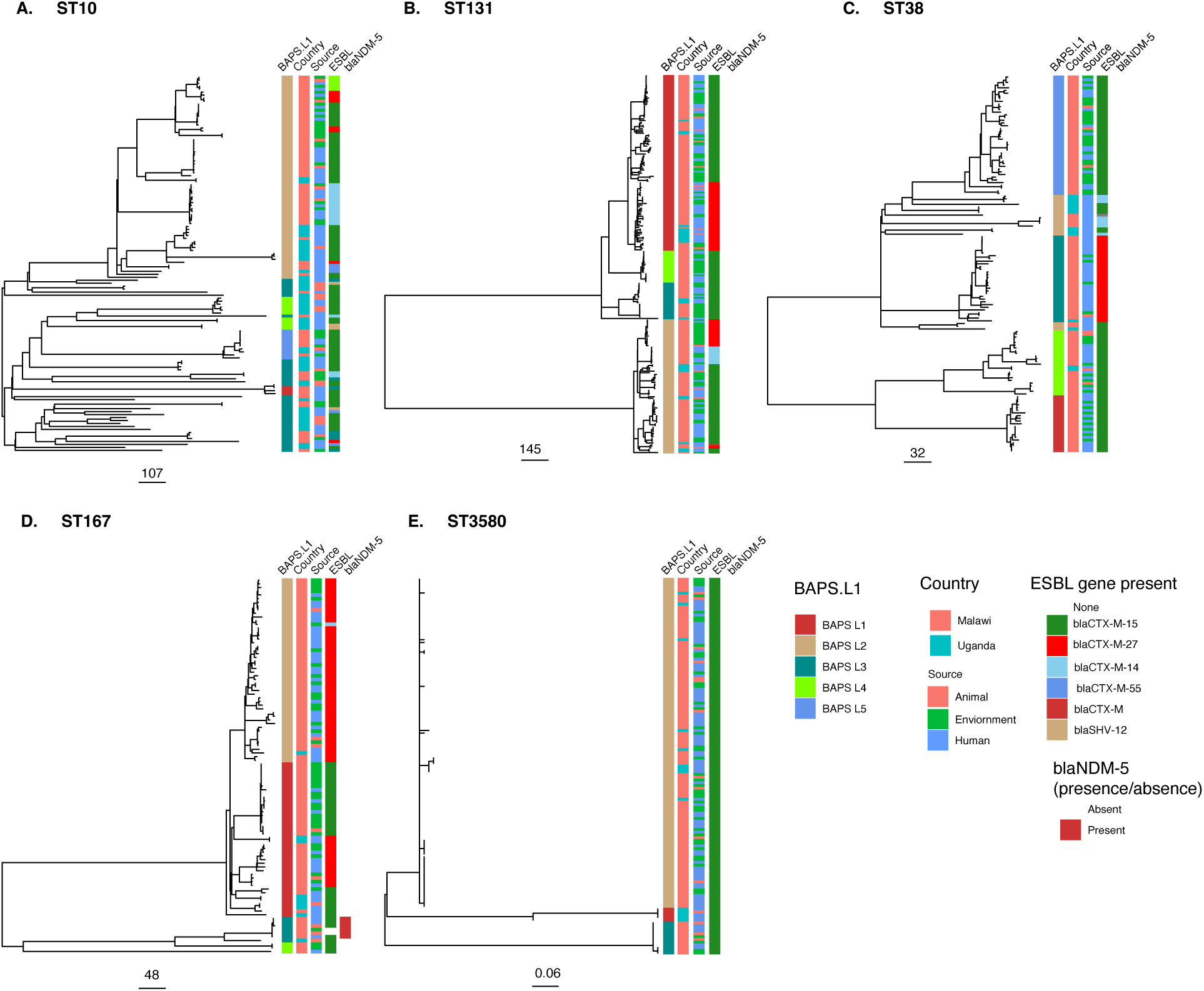
Phylogenetic analysis of common ESBL producing sequence types in Malawi and Uganda indicates overlapping populations between countries and ecological niches. Maximum likelihood phylogenetic trees of (A) ST10 (B) ST131 (C) ST38, (D) ST167 and ST3580 genomes (all n≥ 100) from the DRUM collection. Mapped to trees are level 1 BAPS genome clusters (GC) as identified by hierBAPS, isolate country of origin, sample ecological source, type of ESBL gene present in a genome, and presence or absence of the carbapenemase bla_NDM-5_ gene.

### Inference of transmission events within and between ecological compartments

The phylogenetic intermixing of isolates from different ecological compartments observed in Figure 2, is indicative of sharing or transmission of ESBL-Ec between them. Therefore, to infer putative transmission events, we constructed networks connecting genome pairs at SNP distance thresholds of 0, 1, 2, 5, 10 and 20. We did this for both pairwise core-SNP distances (comprising all 2,344 genomes), and SNP distances inferred from ST-specific reference-based alignments. We found evidence of both within- and between-niche transmission using both the core gene network (Figure 3A) and the ST specific networks (Figure 3 B & C). Notably, we found that despite the human compartment being the most sampled (followed by the environment), human-to-human transmission events were consistently fewer than environment-human transmission events (after deduplication for samples from the same individual across all the SNP thresholds) (Figure 3D). Environmental samples were then re-analysed using specific specimen types, demonstrating that at the highest level of genomic similarity (0-SNP threshold), the most frequent between-niche transmission events were those between human and household water, and human and animal stool (Figure 3E). By comparison, food represented only a very small fraction of inferred transmissions. We overlayed household data to the core SNP network and discovered that isolates from the same household clustered together (Supplementary figure 4). Most core-SNP network clusters however, contained isolates from a mixture of households (Supplementary figure 4), providing evidence for how the within household direct transmission is contributing to the widespread dissemination of ESBL-Ec across communities.

**Figure 3.**
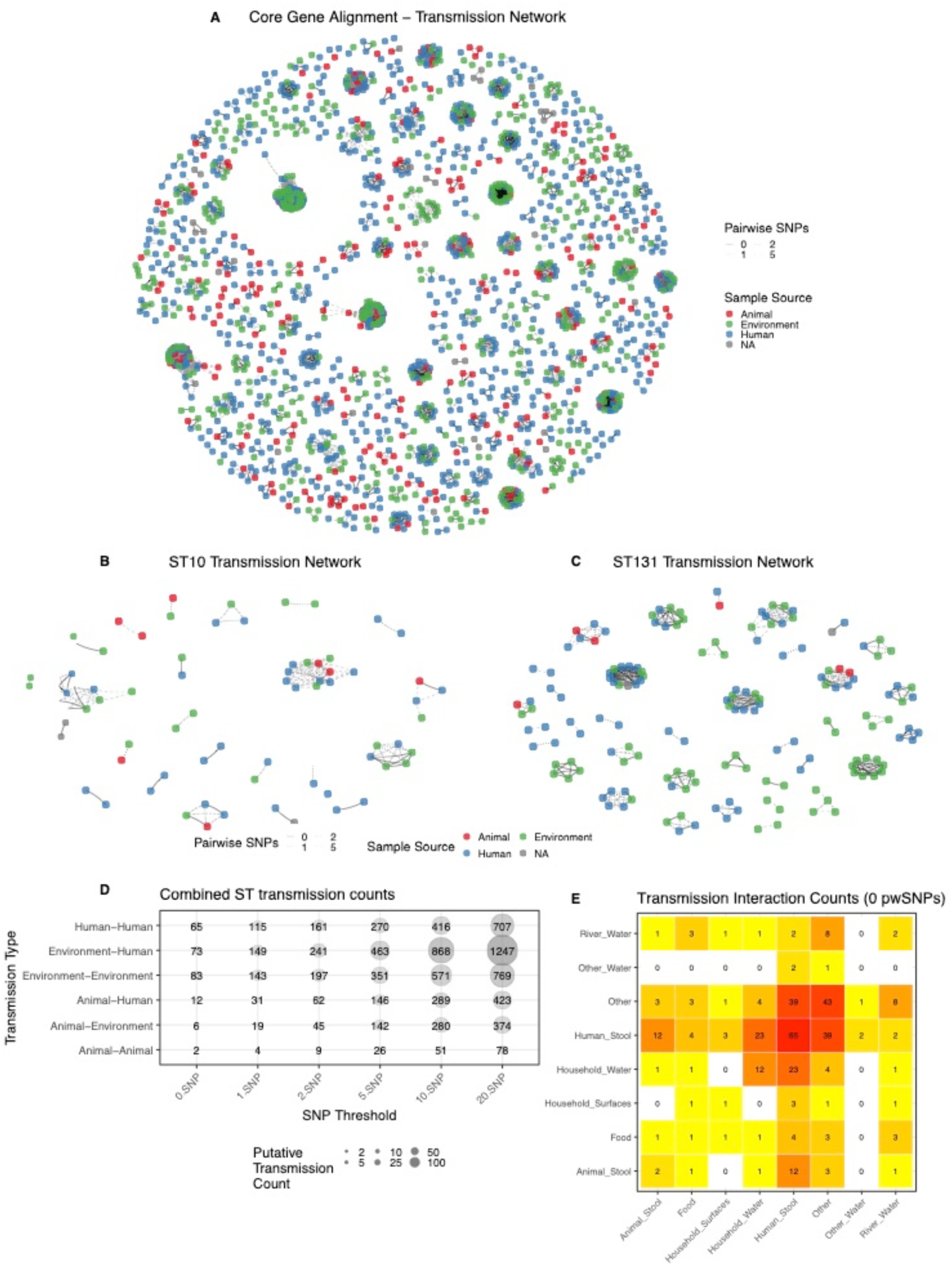
Extremely close genomic relationships between samples from ecological compartments indicates recent transmission from common sources. (A) Core genome pairwise SNP distance-based transmission network for all ESBL E. coli genomes in the DRUM collection with ≤ 5 pairwise SNP distances. Nodes are coloured by source, and edges indicate samples are connected by ≤5 SNPs. Lineage-specific reference mapped analysis of within and between ecological niche transmission networks of ST10 (B) and ST131 (C) genomes with ≤ 5 SNP distances, respectively. (D-E) Counts of transmission events between and within ecological compartments at SNP thresholds of 0, 1, 2, 5, 10 and 20. For panel E, environmental compartment has been split into specific sources of the environmental samples.

### Diversity and distribution of ESBL determinants and associated plasmids

Next, to understand the significance of AMR to our dataset, we screened for and identified 150 unique genes and point mutations across all the 2,344 genomes known to confer resistance to antimicrobial agents of different classes (Supplementary figure 5). Most of these genes or mutations were present in isolates from all the three ecological compartments (human stool, animal stool and the environment) (Supplementary figure 5). We identified 20 ESBL genes; *bla*_CTX-M-15_ (1604/2344 [68.4%], was most common, followed by *bla*_CTX-M-27_ (336/2344 [14.%]) and *bla*_CTX-M-14_ (143/2344 [6.1%]), with other ESBL genes less frequently distributed across genomes (Figure 3A). The distribution of ESBL genes varied across different STs, with some STs such as ST10, ST131 (Figure 2A-D), having more than one ESBL gene, while isolates in STs such as ST3580 (Figure 2E) carried only one ESBL gene. Where more than one ESBL gene was present in a single ST, there was evidence of genes being linked to specific phylogenetic clusters (Figure 2A-E) Additionally, we identified the carbapenamase gene *bla*_NDM-5_ in six genomes and colistin resistance associated *pmrB*_E123D and *pmrB*_Y358N mutations in 14.6% 343/2344 and 28.8% (674/2344) of genomes, respectively (Supplementary figure 5). The chromosomal mutations *pmrB*_E123D and *pmrB*_Y358N were spread across multiple lineages with isolates from all the three ecological niches (Supplementary figure 5). This is indicative of convergent evolution as the pathway for the dissemination of this colistin resistance mechanism. On the other hand, the six genomes carrying the *bla*_NDM-5_ all belonged to isolates from a distinct genome cluster of ST167, of which three were from human, two from animal and one from environmental sources (Figure 2D & Supplementary figure 5). These six isolates came from one household, suggesting a common source.

Given plasmids are one of the primary vectors of AMR genes, we reconstructed and clustered individual plasmid sequences from our genome assemblies using MOB-suite to understand their diversity across compartments in this setting and their role in the distribution of AMR genes, with a focus on ESBLs. This revealed a high diversity of plasmids shared across compartments with 595 distinct plasmid clusters overall, of which 193/595 (32.4%) carried AMR genes, and 89/595 (15.0%) clusters carried ESBL genes. Figure 4C clearly demonstrates the importance of studying both AMR gene and vector independently: individual ESBL genes were widely distributed across multiple plasmid clusters for example *bla*_CTX-M-15_ was found in 55 plasmid clusters, *bla*_CTX-M-27_ in 30 plasmid clusters and *bla*_CTX-M-14_ in 21 plasmid clusters. Conversely, the majority of plasmid clusters were linked to one (48/89 [53.5%]) or two (22/89[24.7%]) ESBL genes. This finding includes the dominant plasmid clusters found in both countries AC315, AA324 and AA887, linked exclusively to *bla*_CTX-M-15_ (Figure 4C). The maximum number of distinct ESBL genes seen in an individual plasmid cluster was eight (Cluster AA474); this observation was also true for other (non-ESBL) AMR genes (Supplementary figure 6). Fitting a fixed effects multinomial regression to the plasmid cluster data with country and ecological source as predictor variables, we found that most plasmid clusters were not associated with a particular country. Exceptions being four and five plasmid clusters less likely to be identified in isolates from Uganda relative to Malawi (AA172, AA738, AA749 and AD094) or more likely to be identified in isolates from Uganda than Malawi (AA282, AA347, A474, AA747 and AC315), respectively (Supplementary figure 7). In addition, cluster AA747 was more likely to be identified in environmental isolates and AD094 more likely to be identified with environmental and human samples than animal isolates (Supplementary figure 7). No significant association was seen for the remaining 80 plasmid clusters with either country of origin or ecological source (Supplementary figure 6).

**Figure 4.**
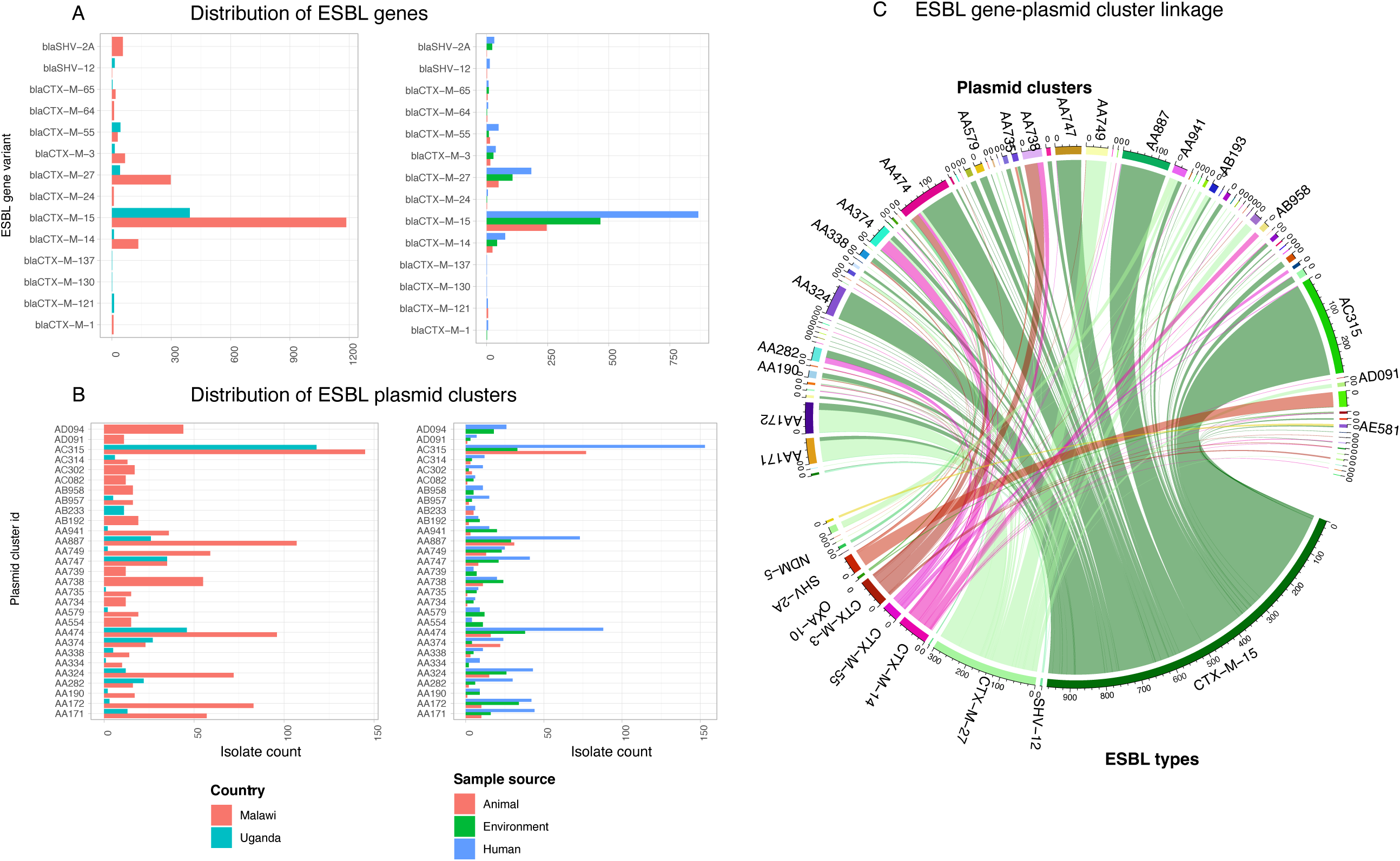
ESBL genes and plasmids are highly diverse and overlap across ecological compartments and countries. A. Distribution of ESBL gene variants by country and ecological compartment (B) Distribution of ESBL gene-carrying plasmid clusters by country and ecological compartment. (C) Circos plot showing links between plasmids clusters (top tracks) and the ESBL type (bottom track) encoded by the gene encoded on the plasmid.

Finally, we overlayed the plasmid cluster data to the core-SNP bacterial host transmission network (Figure 5). Although there were core-SNP clusters with a mixture of plasmids, isolates in most core-SNP network clusters shared the same ESBL plasmid, meaning that the isolates in these clusters were not only similar at chromosomal level, but also shared the same ESBL plasmids. This provides further of evidence of transmission between the different ecological compartments. However, some of the common plasmid clusters such as AC315, AA474 and AA887 were shared across multiple bacterial host core-SNP clusters (Figure 5). We also identified a substantial number of clusters comprising of genomes with ESBL genes but not linked to any plasmid cluster. We investigated the genomic environments of the ESBL genes in some of the genomes found that some of the ESBL genes were on a chromosome (Supplementary figure 8). Genomes not linked to any ESBL plasmid included 639/1,604 (39.8%) genomes carrying *bla*_CTX-M-15_, which is known to be both plasmid- and chromosome-borne.

**Figure 5.**
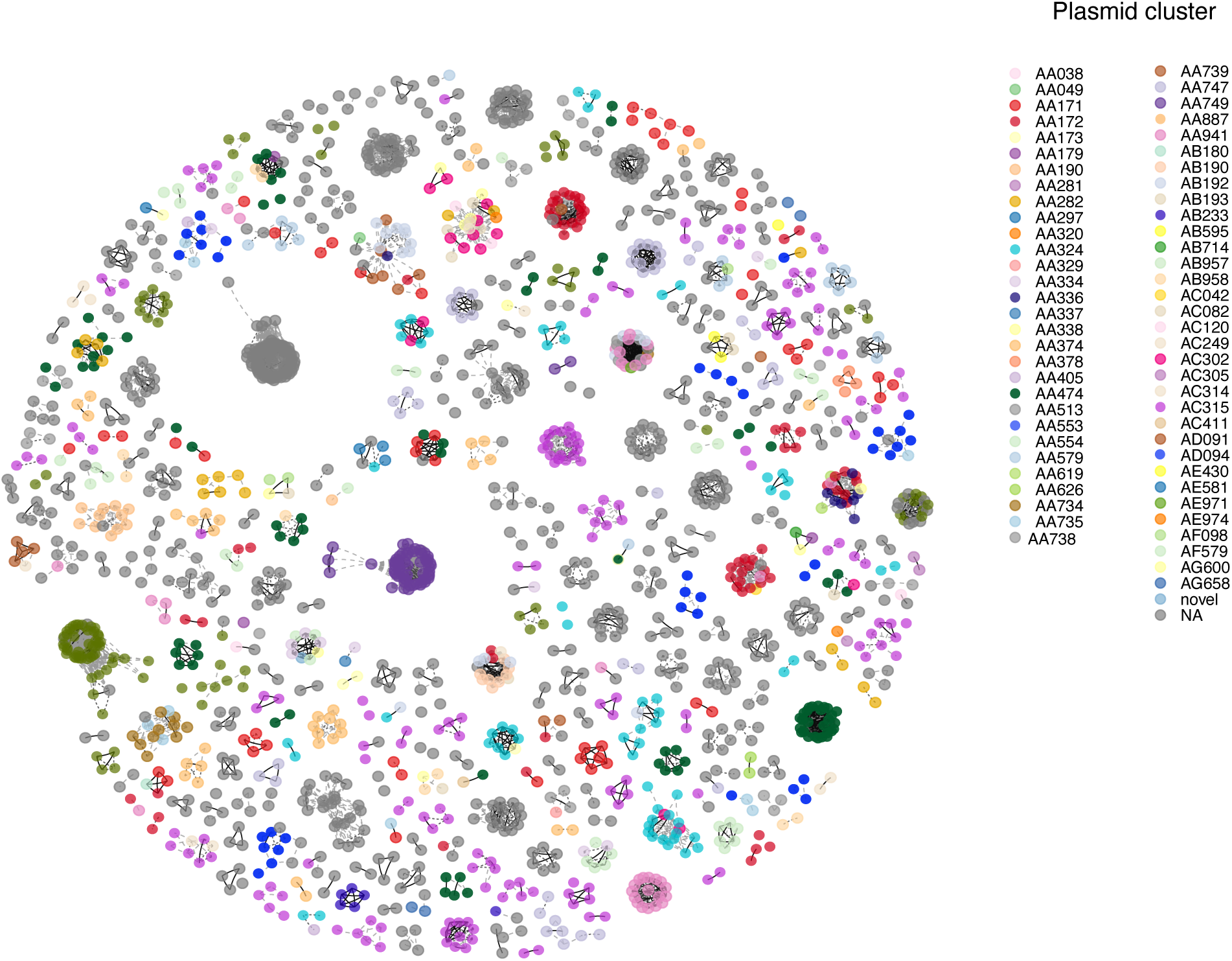
Inferred transmission clusters can carry the same or different ESBL plasmid. Core genome pairwise SNP distance-based transmission network for all ESBL E. coli genomes in the DRUM collection with ≤ 5 pairwise SNP distances. Nodes represent genomes coloured by ESBL-plasmid cluster (pale grey genome nodes lack an ESBL plasmid), and edges indicate sample connections by ≤5 SNPs. Clusters of linked genomes form putative transmission networks: the majority of genomes in these putative networks carry the same ESBL plasmid cluster, although some mixing is also visible within clusters.

## Discussion

In this study, we investigated the genomic diversity and transmission of ESBL-Ec in Malawi and Uganda using prospective, longitudinally collected samples from humans, animals and the environment. All samples were epidemiologically linked at the household level, or in the case of river water, the geographic area in which the study was being conducted.^16^ The direct epidemiological link between genetic diversity to ecological source enabled us to unravel ongoing transmission between these ecological compartments, both within and between households. Here, we showed that ESBL-Ec were highly diverse. This high diversity was observed at both the strain level and also at the level of the ESBL genetic determinants (including both the ESBL genes themselves, and the plasmids which harbour them). However, despite the huge diversity, we found clusters of closely related ESBL-Ec strains and ESBL determinants that were shared by different ecological compartments, indicating frequent sharing or transmission of ESBL-Ec and/or ESBL determinants between these compartments. While there were differences in distribution of isolates by lineage between Malawi and Uganda, most STs and all the ESBL genes and plasmids were found in both countries indicating that the transmission of these strains and the ESBL gene flow is further not limited by national borders.

Globally, ESBL production in *E. coli* has been associated with a limited number of STs and AMR genes, with ST131 and *bla*_CTX-M-15_ being the most prominent.^17,18^ This is true for Malawi where ST131 and *bla*_CTX-M-15_ have previously been identified as the leading ESBL ST and gene respectively. However, this study has captured far more strain and ESBL gene diversity than previous studies.^19,20^ For Uganda, we found a somewhat contrasting genomic epidemiology of ESBL-Ec, in that ST10 rather than ST131 was the dominant ESBL-Ec. The high diversity of ESBL-Ec that this study has captured in both Malawi and Uganda demonstrates the widespread dissemination of ESBL determinants, and in particular of *bla*_CTXM-15_ to lineages they were not previously observed in, such as the novel STs identified through Enterobase (Supplementary Table 1). Whilst the high burden of bacterial infections and subsequent high usage of 3GC in SSA should be expected to drive the acquisition and spread of ESBL-Ec, the sharing of plasmids by distinct multiple lineages and transmission clusters means that even when transmission cannot occur at a bacterial host strain level, plasmids facilitate horizontal transfer of ESBL genes across lineages and the genes between plasmid vectors. Given the nature and likely frequency of exposure and transmission we have revealed in this setting, our data describes a truly One Health view of AMR at the level of: the bacterial host, the plasmid vector and the individual AMR gene and across ecological compartments. It did not escape our notice that while the focus of this study was on ESBL determinants, in addition to the ESBL genes, we also found genes encoding resistance to reserve antimicrobials including carbapenems (*bla*_NDM-5_) and colistin in isolates from humans, animals and the environment. Carbapenem use in Malawi is limited but growing, whereas although there is no recorded use of colistin in humans and its use is banned in animals in Malawi, a recent study revealed that colistin is one of the antimicrobial agents commonly dispensed from veterinary shops.^22^

To develop effective interventions against AMR transmission, our study underlines the need to use genomic data to understand reservoirs and routes of AMR transmission in different geographical and socio-economic contexts. It provides a unique view of AMR in this setting revealing the multiple pathways for successful dissemination of ESBL genes. More specifically, the reservoirs of AMR genes may not necessarily be clinical nor human, and the One Health framework provides a holistic approach for addressing the AMR problem.^21^ The One Health framework informed our sampling strategy, and we have shown that here highly similar ESBL-Ec strains and ESBL determinants are often shared between humans, animals and the environment. These findings perhaps understandably, contrast markedly that of studies from Europe with high antimicrobial stewardship which suggests limited transmission of AMR bacteria or flow of genetic determinants of AMR between different ecological compartments.^23^ In rural and peri-urban settings such as those of our study, human interactions with animals and the environment are fundamentally different to that of many high-income settings. It is common for domestic animals including poultry, goats and pets such as dogs and cats to share homes with humans.^24^ Specifically, we have identified household stored water to be a major source or reservoir of ESBL-Ec shared with humans. This could be linked to poor water, sanitation and hygiene (WASH) practices and infrastructure, which lead to high levels of enteric bacteria being shed into the environment through human and animal faeces. High frequency of contact between humans, animals and the environment, likely facilitates increased transmission of ESBL-Ec and other enteric bacteria between the different ecological sources. What is unclear however, is the directionality of the transmission between the different ecological compartments.

## Conclusion

By taking a One Health approach to the study of ESBL*-*Ec, we reveal through the lens of genomics that in Eastern Africa, genetically similar ESBL-Ec strains and ESBL determinants are shared and transmitted between all relevant ecological compartments at the level of the individual household and that this occurs at every household we have sampled. The success of different *E. coli* ST’s, the mobility of ESBL genes under selection from high 3GC exposure to occupy multiple plasmid vectors and ability of plasmids to successfully disseminate these genes into new lineages provide multiple pathways for the successful dissemination of ESBL genes across households and between country states in SSA. Using ESBL-Ec as an exemplar, we provide the high resolution data showing AMR is indeed a One Health problem in an African context.

## Methods

### Study isolates

The DRUM household study was conducted from April 2019 to November 2020 at three sites in Malawi including Chileka, Chikwawa and Ndirande; and from July 2020 to August 2021 at two sites in Uganda, including Kampala and Hoima. Ndirande (Blantyre, Malawi) and Kampala (Uganda) represented urban settings, Chileka (Blantyre) a peri-urban setting, whereas Chikwawa represented a rural setting, and Hoima represented both rural and peri-urban settings. Two hundred fity-nine households were enrolled at baseline and followed up at one month, three months and six months in Malawi; an additional ninety-two households were enrolled at baseline and followed up at one month, two months and four months in Uganda. On each household visit, samples collected included human and animal stool, household-linked environmental samples such as food, water, door handles and clothes; and broader environmental samples such as river water. Data collected at each household visit included individual and household demographics, antimicrobial use, health seeking behaviour and WASH behavioural practices. Detailed descriptions of these data and the collection procedures are published elsewhere.^16,24^

### Microbiology

All samples were initially inoculated into enrichment broth (Buffered Peptone Water-BPW) and then placed in an aerobic incubator at 37 ± 1°C for 18–24 hours. After incubation a 1.8 ml aliquot of the culture BPW was stored at -80°C, and a 1µl loop of the remaining sample plated onto ESBL chromogenic agar (CHROMagar™, France). Plates were placed in aerobic incubator at 37 ± 1 °C for 18–24 hrs and then read for growth of ESBL bacteria. Pink colonies and (indole positive) white colonies were categorised as ESBL-Ec.^16^

### DNA extraction, library preparation and sequencing

DNA was extracted from all ESBL-positive single colony isolates using the QIASymphony DSP Virus/Pathogen mini-kit® on the QIASymphony® (QIAGEN, USA) automated DNA extraction platform or manually extracted using the DNeasy® blood and tissue kit (QIAGEN, USA) at Malawi-Liverpool-Wellcome Programme (MLW), Malawi and the Department of Medical Microbiology, Makere University, Uganda. DNA extracts were shipped to Wellcome Sanger Institute (WSI) where DNA libraries were constructed using NEB Ultra II custom kit on an Agilent Bravo WS automation system and sequenced on the Illumina HiSeq X10 platform (Illumina Inc, California, USA) to produce paired-end raw reads of 150 base pairs (bp). Quality of raw reads was assessed using FastQC (https://www.bioinformatics.babraham.ac.uk/projects/fastqc/) and MultiQC.^25^ We used Kraken v0.10.6 to confirm sample species and excluded genomes with < 40% of raw reads belonging to *E. coli*.^26^ Raw sequence data were deposited in the European Nucleotide Archive (ENA) and ENA accession numbers are included in Table S1.

For 38 selected samples identified as representative of major lineages or MLSTs, we reextracted DNA from the colony isolate using the MasterPure kit (BioSearch Technologies, USA) and performed long-read sequencing on Sequel II (PacBio, USA).

### *De novo* assembly and sequence annotation

We ran an WSI automated pipeline to assemble raw reads into contiguous sequences (contigs) using Spades (v3.14.0) and annotated the assemblies using PROKKA (v1.14.5).^27–29^ Quality assessment of genome sequence assemblies was performed as follows: we filtered out contigs of length <300 bp; generated assembly statistics and excluded from further analysis assemblies with total size < 4 mega base pairs (MB) or > 6MB or with N50 < 80,000 bp; We assessed genome completeness and contamination using checkM (v1.2.2); and excluded genomes with <90% completeness or > 5% contamination.^30^

### Multi-locus sequence typing and characterisation of AMR determinants and plasmids

Multi-locus sequence typing (MLST) was performed *in silico* using mlst tool (v2.16.2) (https://github.com/tseemann/mlst). We screened for AMR determinants using AMRfinder plus (v3.10.40) and used the mob-recon function in mob-suite v3.0.3 to identify and cluster plasmid contigs.^31,32^ Plasmid replicon types were assigned using mob-suite and by running abricate (https://github.com/tseemann/abricate) against the PlasmidFinder database.^31,33,34^ Where relevant, we used minimum thresholds of 95% blastp identity and 90% coverage.

### Core genome phylogeny reconstruction and population structure analysis

We inferred a pan-genome using the default settings of panaroo v1.3.0 and classified genes identified in at least 99% of the genomes as core.^35^ We concatenated alignments of the core genes to generate a core gene alignment. Single nucleotide polymorphic (SNP) sites were extracted from the core gene alignment using snp-sites v2.5.1 to generate a core-SNP alignment.^36^ A core gene maximum likelihood (ML) phylogenetic tree was constructed from the core SNP alignment using IQ-TREE v1.6.12 GTR+I+G model.^37^ Reliability of the inferred branches and branch partitions in the phylogenetic tree was assessed with 1000 ultrafast bootstrap replicates using UFBoot2.^38^ Phylogenetic trees were visualised and annotated using ggtree v3.5.1.901 package in R v4.2.1 and iTOL.^39–41^ We performed population structure analysis on the whole collection using PopPUNK v2.6 and assigned lineages based on a previously described PopPUNK reference database for *E. coli.*^42,43^

### Lineage specific analyses

We performed lineage specific analyses on the five most common STs in this collection (ST131, ST38, ST10, ST617 and ST3580), each comprising 100 or more genomes. We selected 2-4 genomes from each of these five STs and performed long-read sequencing using the Pacific Biosciences (PacBio) platform. We assembled *de novo* the resulting long-read raw data with flye v2.9.2 assembler and polished the assemblies with their corresponding Illumina short-reads using polypolish v0.5.0.^44,45^ We used Quast v5.0.2 to assess the quality of the generated hybrid assemblies.^46^ For each of the four STs, the hybrid assembly with the highest quality scores was selected as the local ST-specific reference genome, to which all short reads of genomes in that ST were mapped using BWA aligner followed by INDEL realignment using GATK v3.4.46 Indel Realigner.^47,48^ Variant calling and consensus pseudosequence generation were performed using samtools v1.2 and bcftools v1.2. A minimum of 8 supporting reads (3 per strand) and a variant frequency/mapping quality cut-off of 0.8 were used to call variants. Sites not meeting these criteria were masked to ‘N’ in the pseudosequence. We used Gubbins v3.2.1 to identify and mask recombination sites and constructed recombination-free phylogenetic trees with IQTree.^49^ We performed ST-specific population structure analyses using RhierBAPS.^50 18^

### SNP Analysis

For ST-specific multiple sequence alignments (for ST10, ST38, ST131, ST167), we inferred pairwise SNP distances using pairsnp v0.1.0 (available at https://github.com/gtonkinhill/pairsnp). We linked metadata to samples in R and evaluated the distribution of pairwise SNP distances for each ST sequence alignment within and between countries, households, and individual people, examining SNP distributions to determine a threshold for including the 5% closest comparisons (≤2 SNPs in all cases). We inferred putative transmission networks by constructing edge-networks from all pairwise comparisons below a series of thresholds (0, 1, 2, 5, 10 and 20 SNPs) using the network v1.18.1 and iGraph v1.5.0 packages, and these were plotted using ggnetwork v0.5.12 in R^51–53^. We quantified putative transmission events at each threshold by counting the number of pairwise comparisons between each sample source type (e.g. human, environment, animal). Where multiple samples from each human individual were present, we deduplicated pairwise comparisons less than or equal to the threshold being evaluated for transmission.

### Statistics

Associations between ST versus country and ST versus ecological compartment were performed using chi-square test or Fisher’s exact test where appropriate. We performed multinomial regression analysis to determine the associations between plasmid clusters and sample country of origin and ecological source using the nnet package in R.

### Ethical approval

This study was approved by the College of Medicine Research Ethics Committee in Malawi (No. P.11/18/2541), Makerere University School of Veterinary Medicine and Animal Resources REC (Ref: SVARREC/18/2018) (Ref: SVARREC/18/2018), Joint Clinical Research Centre (JCRC) REC, (No. JC3818) and Uganda National Council for Science and Technology (UNCST, No. HS489ES) in Uganda and LSTM Research and Ethics Committee (REC, No. 18-090) in United Kingdom. Transfers of the bacterial DNA from MLW and IDI to the Wellcome Sanger Institute were compliant with the Nagoya Protocol on Access and Benefits Sharing under export permit number MEPA-12-07-1313-21-16c in Malawi and waived by the UNCST in Uganda as research intended for educational purposes.

## Data availability

Raw sequence data were deposited in the European Nucleotide Archive under project ID PRJEB40384.

## Acknowledgements

We would like to thank the study participants in Malawi and Uganda, the DRUM field and laboratory teams at Malawi-Liverpool-Wellcome Programme (MLW) and the Infectious Disease Institute (IDI), Makerere University. We also thank the data teams at MLW, IDI and Lancaster University and members of <u>The DRUM Consortium</u>. We acknowledge the Core Sequencing and Pathogen Informatics teams at the Wellcome Sanger Institute.

This work was funded by the Antimicrobial Resistance Cross-Council Initiative through a grant from the Medical Research Council, a Council of UK Research and Innovation, and the National Institute for Health Research. P.M. is supported by a Wellcome International Training Fellowship 223012/Z/21/Z. MLW is supported by a Wellcome Trust core grant 206545/Z/17/Z. M.A.B. and N.R.T. are supported by Wellcome funding to the Sanger Institute (no. 206194).

## Author contributions

P.M., N.A.F. and N.R.T. designed the study. D.C., F.O., K.C., G.K., S.N., F.I., L.M., H.K., S.J. and N.A.F. collected the samples and metadata in this study. D.C., F.O., A.Z., and C.S. performed sample processing, microbiology culture and DNA extractions. P.M. and M.A.B. performed the bioinformatics and statistical analyses with guidance from NRT. P.M., M.A.B., N.A.F. and N.R.T. wrote the manuscript. All authors reviewed and commented on the manuscript.

## Competing Interests

The authors declare no competing interests.

## Notes

### Competing Interest Statement

The authors have declared no competing interest.

