## Supplementary data for "One Health in Eastern Africa: No barriers for ESBL producing *E. coli* transmission or independent antimicrobial resistance gene flow across ecological compartments"

AUTHOR AFFILITATION:

[1]Wellcome Sanger Institute, Wellcome Genome Campus, Hinxton, Cambridgeshire, UK;

[2]Malawi-Liverpool-Wellcome Programme, Kamuzu University of Health Sciences, Blantyre, Malawi;

[3]Department of Vector Biology, Liverpool School of Tropical Medicine, Liverpool, UK;

[4]Department of Clinical Sciences, Liverpool School of Tropical Medicine, Liverpool, UK;

[5]Infectious Disease Institute, Makerere University, Kampala, Uganda;

[6]College of Health Sciences, Makerere University, Kampala, Uganda;

[7] Malawi University of Business and Applied Sciences, Blantyre, Malawi

[8]College of Veterinary Medicine, Animal Resources and Biosecurity, Makerere University, Kampala, Uganda;

[9]School of Public Health, Makerere University, Kampala, Uganda;

[10]Department of Civil and Environmental Engineering, University of Strathclyde, Glasgow, UK;

[11]WALIMU, Kampala, Uganda;

[12]School of Medicine, St Andrews University, St Andrews, Fife, UK;

[13]London School of Hygiene and Tropical Medicine, London, UK.

**# Corresponding Authors:**

Patrick Musicha:

Nicholas R Thomson:

### Supplementary material

#### Supplementary figures

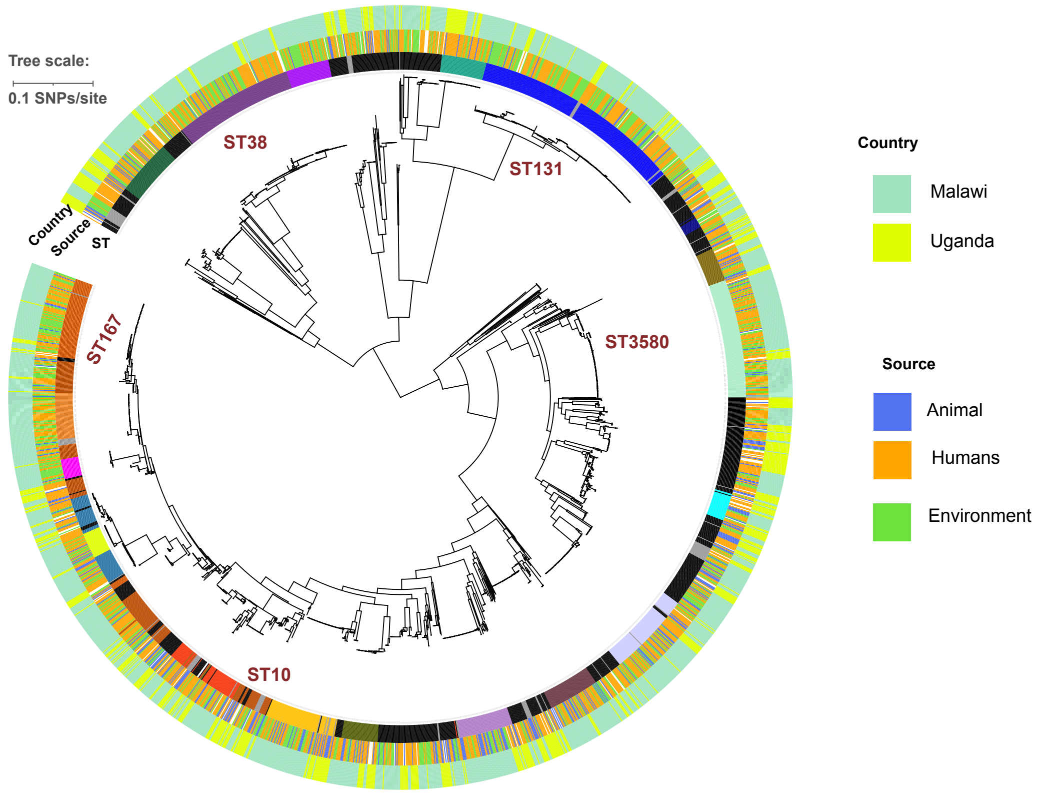


***Supplementary figure 1:* Maximum likelihood core-genome phylogenetic tree of ESBL producing carriage E. coli from Uganda and Malawi.** The phylogenetic tree is mid-point rooted. The inner ring shows sequence type distribution across the tree, the middle ring shows isolate ecological source as human stool, animal stool or the environment and the outer ring shows isolate country of origin, either Malawi or Uganda.

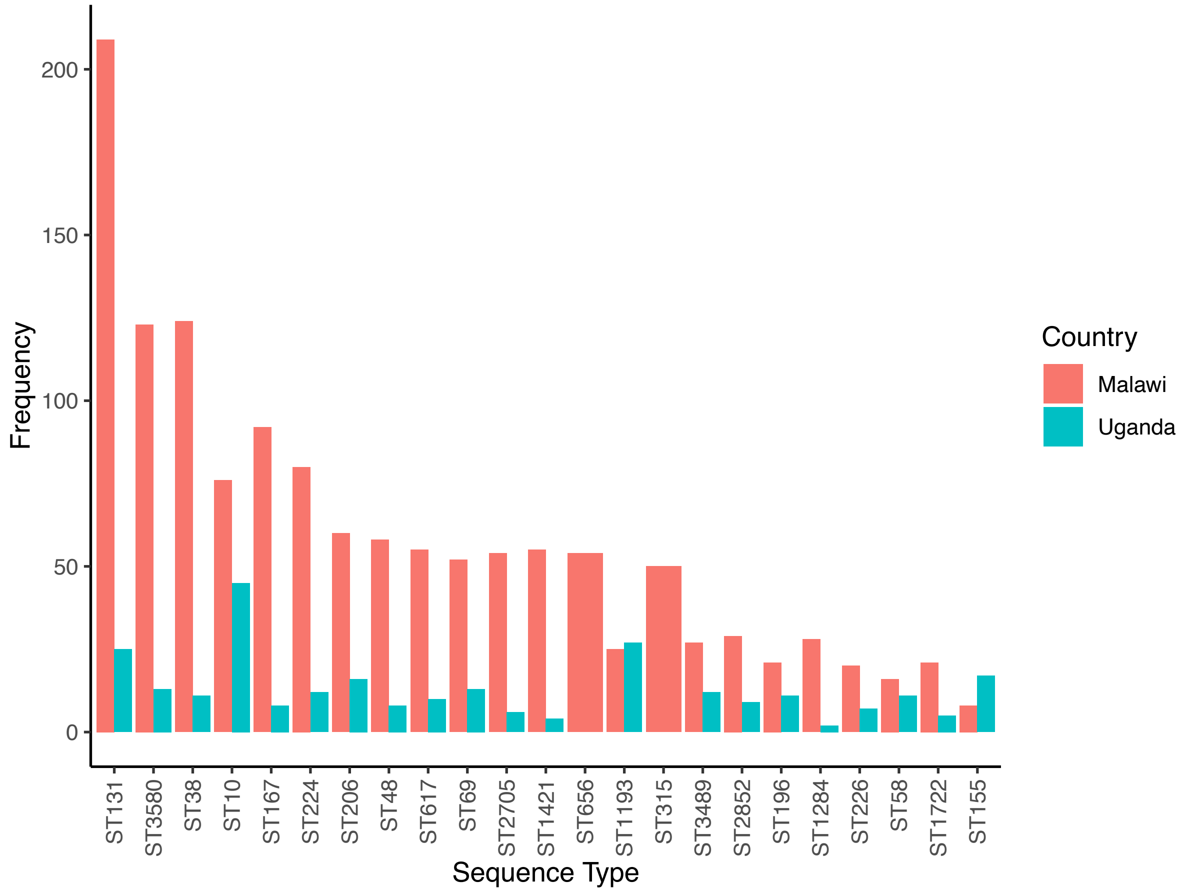


***Supplementary figure 2*: Distribution of sequence types with at least 1% of genomes, by sample country of origin**

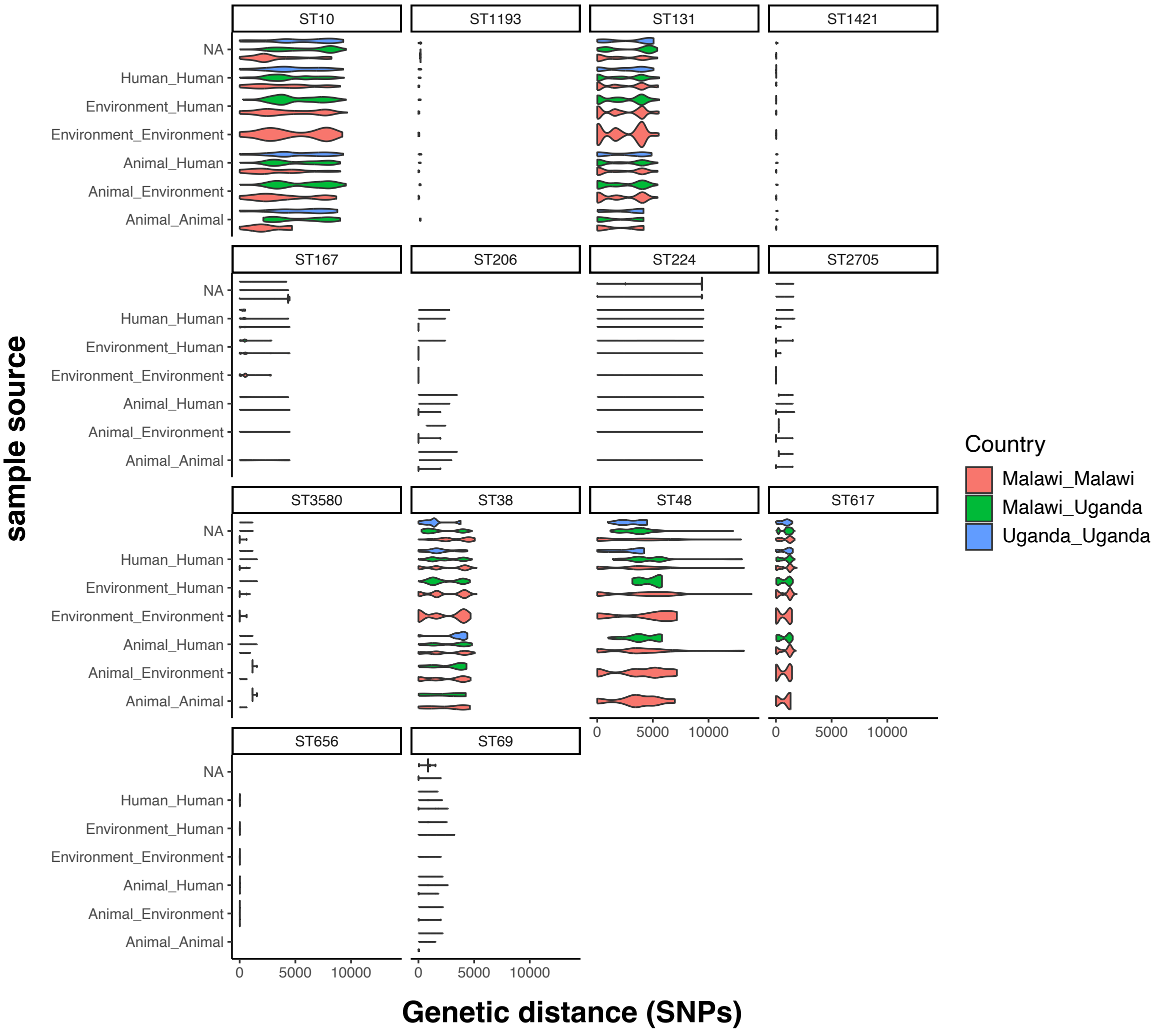


***Supplementary figure 3*: Distribution of core genome SNP distances**

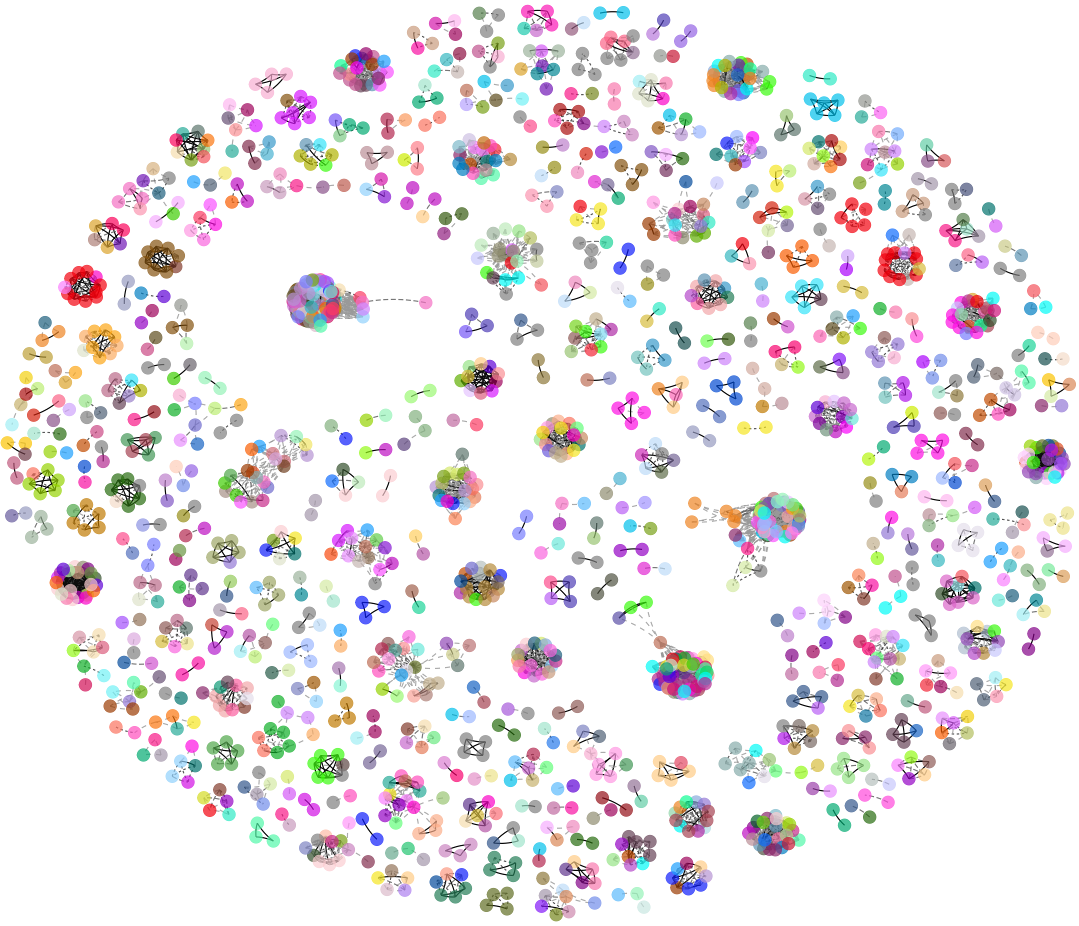

***Supplementary figure 4:*** **ESBL E. coli Transmission cluster-household linkage.** A core genome pairwise SNP distance-based transmission network for all ESBL E. coli genomes in the DRUM collection with ≤ 5 pairwise SNP distances coloured. Each network node represents a genome and is coloured by the sample household ID.

Extended data.

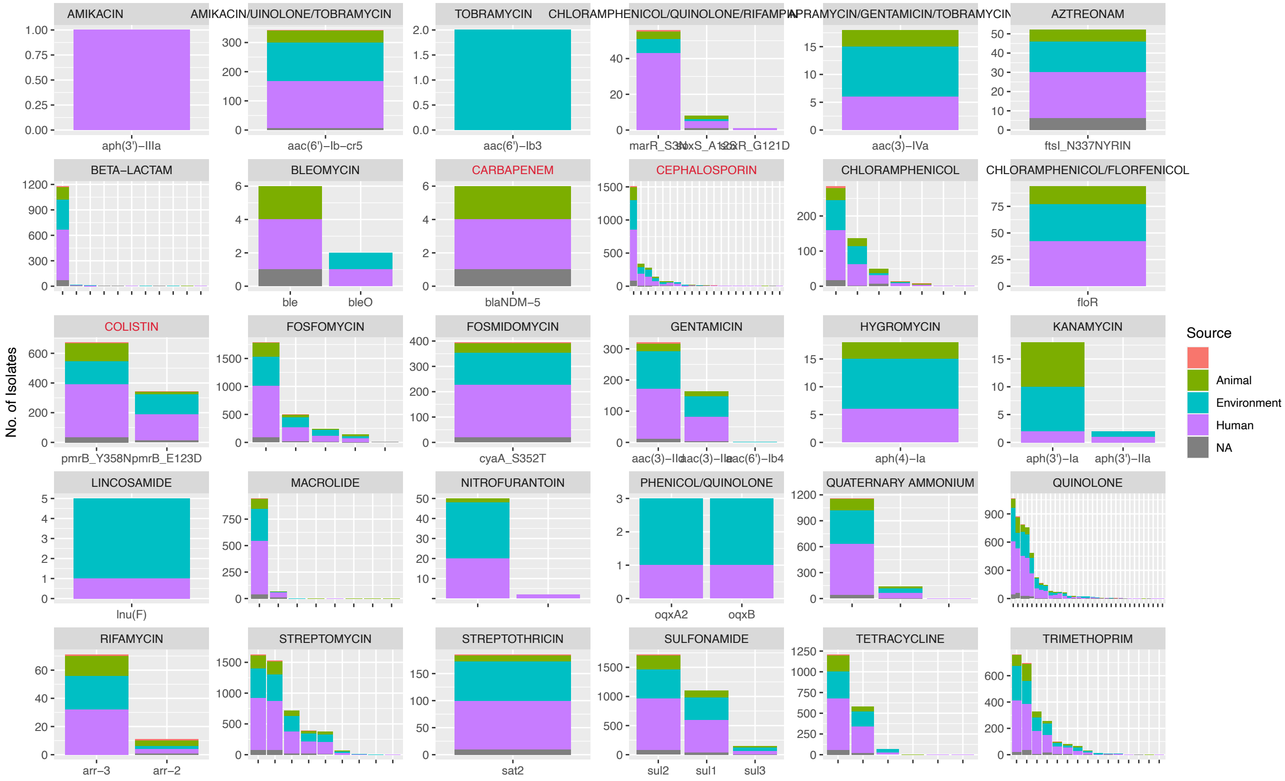


***Supplementary figure 5:*** **Distribution of AMR genes.** Each bar plot shows the distribution of AMR genes for a given class of antimicrobials by sample ecological source.

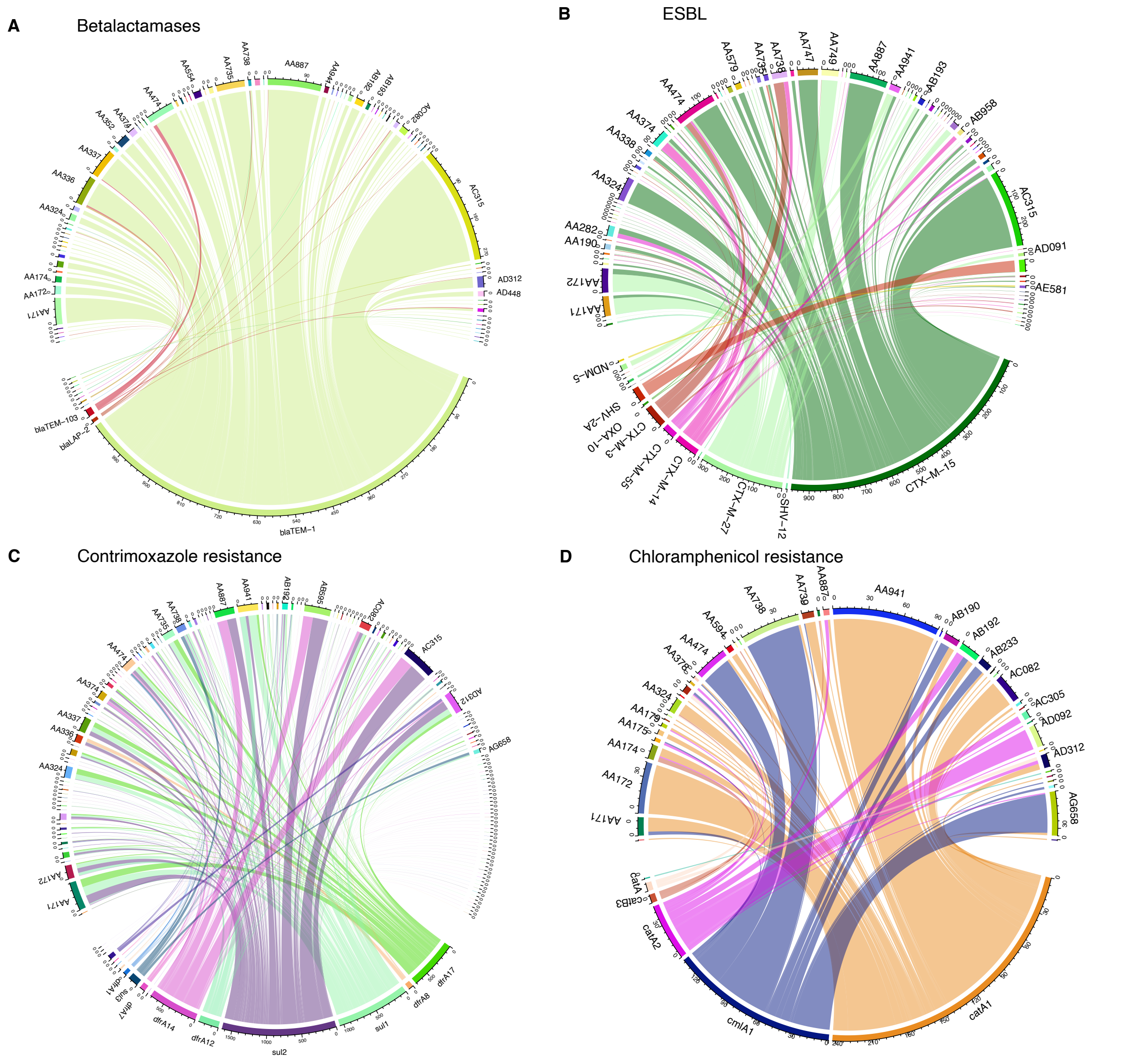


***Supplementary figure 6:*** **Circos plots showing links between plasmids clusters and genes they carry encoding resistance to (a) narrow spectrum beta-lactams (b) 3rd generation cephalosporins (extended spectrum beta-lactams (ESBLs)) (c) cotrimoxazole and (d) chloramphenicol .**

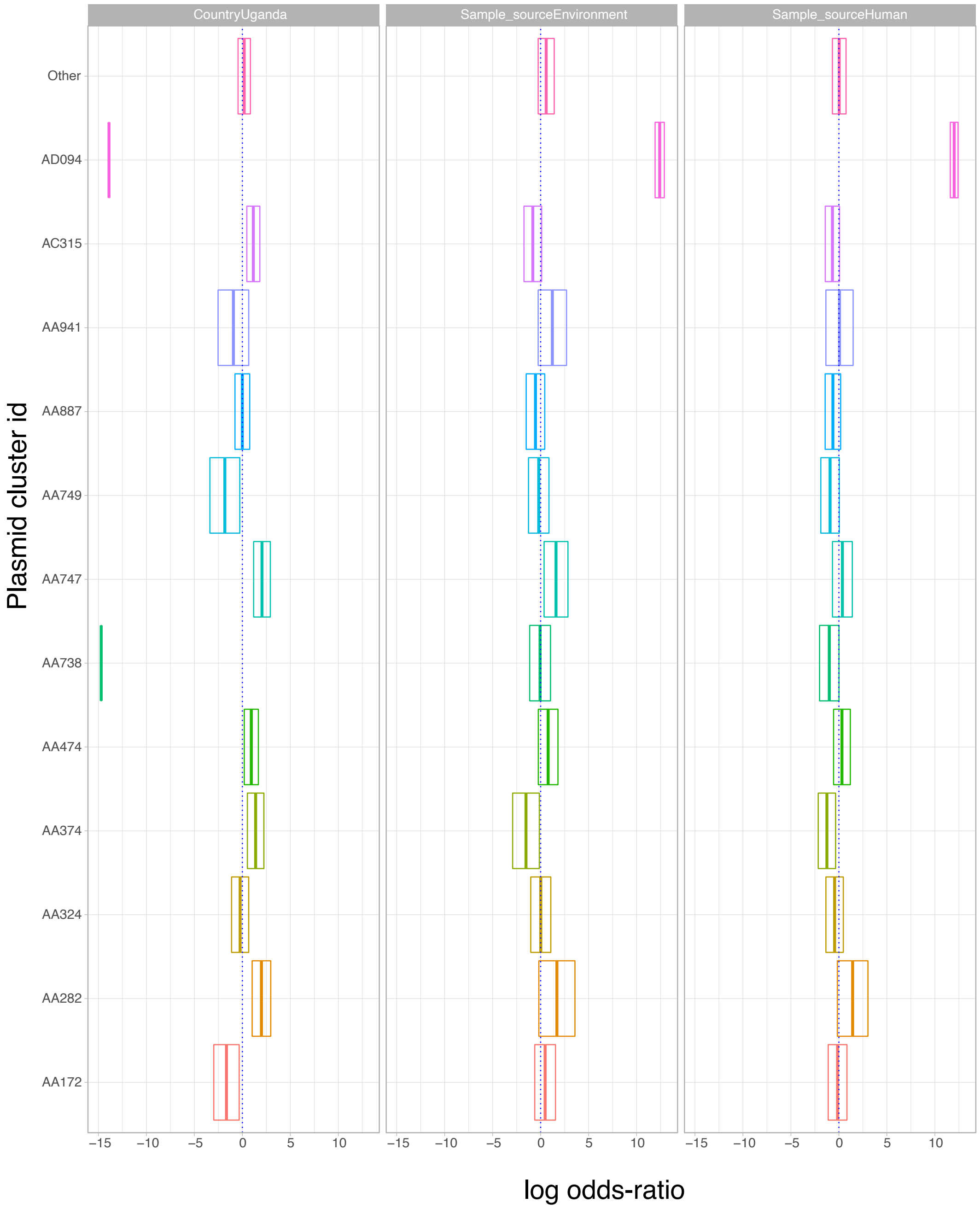


***Supplementary figure 7:* Association between ESBL plasmid and sample country or ecological source.** Associations are presented as cross bars showing odds ratio (on a log scale) of a plasmid cluster being associated with a particular country (Uganda relative to Malawi) or ecological source (environment or human relative to animal). The log-odd ratio values in shown by the middle vertical line of the cross bar and boundaries of the cross bar on the left and right of the odds-ratio line represent the lower and upper bounds of a 95% confidence interval.

**A**

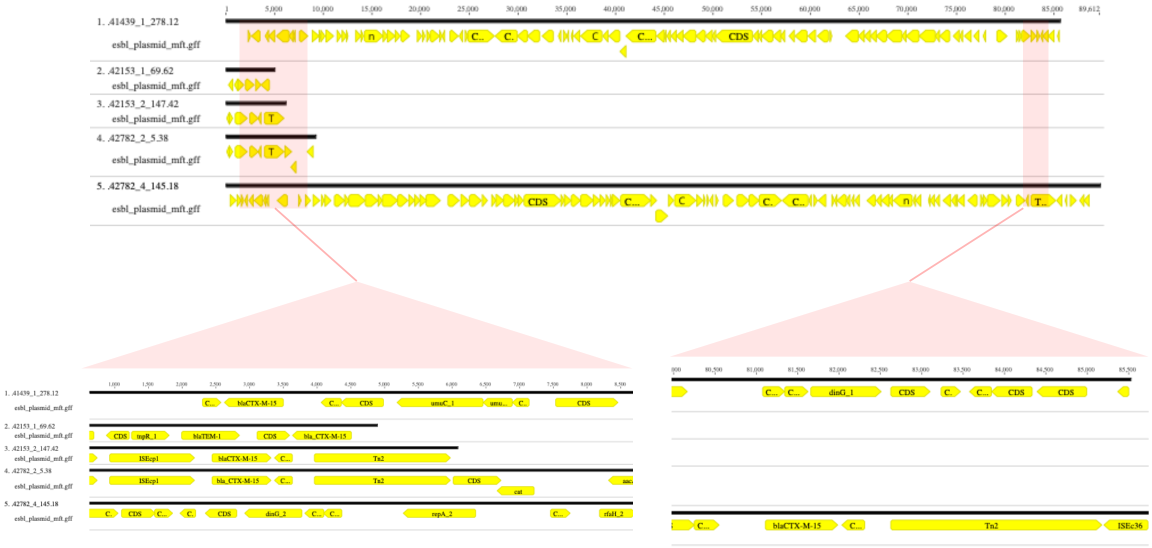

**B**

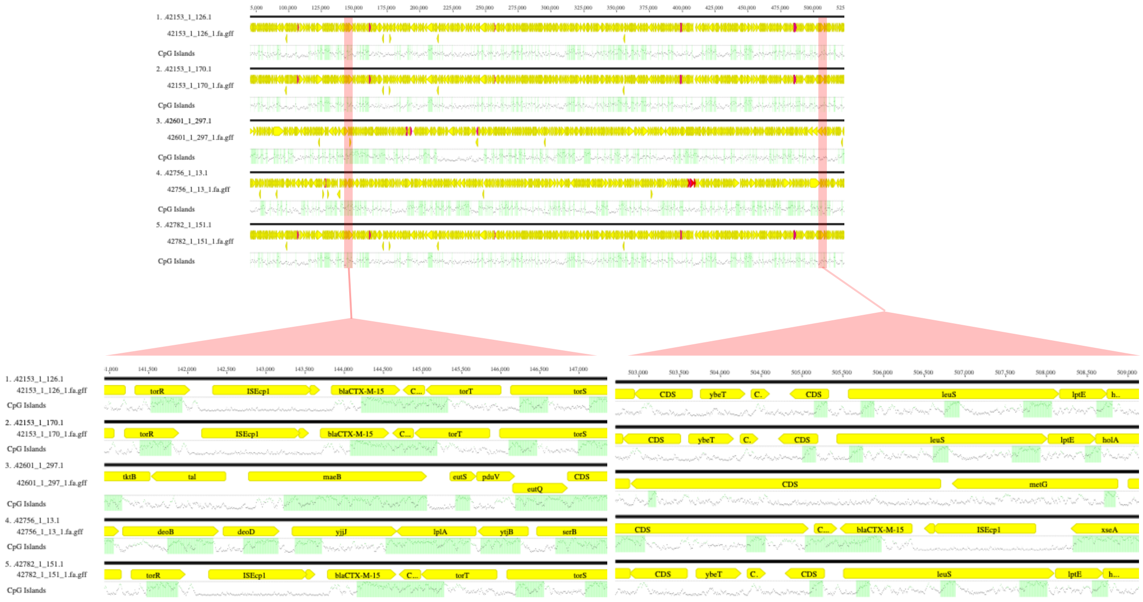

***Supplementary figure 8:*** **Genetic environments of ESBL genes (A) on plasmid and (B) on chromosome sequences**

#### Supplementary tables

***Supplementary table 1:*** **List of isolates submitted to Enterobase for multi-locus sequencing typing and their identified sequence types**

| Sanger_lane_ | Date Entered | Release Date | ST | ST Complex | Lineage | adk | fumC | gyrB | icd | mdh | purA | recA |
|---|---|---|---|---|---|---|---|---|---|---|---|---|
| 41439_143 | 13/12/2023 | 13/12/2023 | 7696 | ST10 Cplx | A | 744 | 11 | 4 | 8 | 8 | 8 | 2 |
| 42601_1_342 | 14/12/2023 | 14/12/2023 | 8025 | ST10 Cplx | A | 10 | 7 | 4 | 8 | 8 | 5 | 2 |
| 42782_1_53 | 08/01/2024 | 08/01/2024 | 8025 | ST10 Cplx | A | 10 | 7 | 4 | 8 | 8 | 5 | 2 |
| 42782_1_147 | 08/01/2024 | 08/01/2024 | 8025 | ST10 Cplx | A | 10 | 7 | 4 | 8 | 8 | 5 | 2 |
| 42782_1_157 | 08/01/2024 | 08/01/2024 | 8025 | ST10 Cplx | A | 10 | 7 | 4 | 8 | 8 | 5 | 2 |
| 42782_1_303 | 08/01/2024 | 08/01/2024 | 8025 | ST10 Cplx | A | 10 | 7 | 4 | 8 | 8 | 5 | 2 |
| 42782_1_328 | 08/01/2024 | 08/01/2024 | 8025 | ST10 Cplx | A | 10 | 7 | 4 | 8 | 8 | 5 | 2 |
| 42782_2_14 | 08/01/2024 | 08/01/2024 | 8025 | ST10 Cplx | A | 10 | 7 | 4 | 8 | 8 | 5 | 2 |
| 41439_1_288 | 13/12/2023 | 13/12/2023 | 8986 | ST10 Cplx | A | 10 | 7 | 4 | 8 | 12 | 109 | 2 |
| 42756_4_180 | 15/12/2023 | 15/12/2023 | 8986 | ST10 Cplx | A | 10 | 7 | 4 | 8 | 12 | 109 | 2 |
| 41330_8_58 | 13/12/2023 | 13/12/2023 | 9525 | ST10 Cplx | A | 10 | 11 | 4 | 140 | 8 | 1 | 2 |
| 41330_8_51 | 13/12/2023 | 13/12/2023 | 13391 | ST10 Cplx | A | 10 | 11 | 1209 | 8 | 8 | 8 | 2 |
| 41439_1_268 | 13/12/2023 | 13/12/2023 | 13391 | ST10 Cplx | A | 10 | 11 | 1209 | 8 | 8 | 8 | 2 |
| 42756_3_126 | 15/12/2023 | 15/12/2023 | 13391 | ST10 Cplx | A | 10 | 11 | 1209 | 8 | 8 | 8 | 2 |
| 42782_4_291 | 10/01/2024 | 10/01/2024 | 13814 | ST10 Cplx | | 6 | 11 | 4 | 1 | 8 | 8 | 133 |
| 42756_3_90 | 15/12/2023 | 15/12/2023 | 14131 | ST10 Cplx | A | 10 | 11 | 4 | 8 | 8 | 1183 | 73 |
| 41439_1_13 | 13/12/2023 | 13/12/2023 | 14381 | ST10 Cplx | | 6 | 11 | 1441 | 8 | 8 | 8 | 2 |
| 41439_1_24 | 13/12/2023 | 13/12/2023 | 15381 | ST10 Cplx | A | 6 | 2452 | 4 | 8 | 8 | 8 | 2 |
| 41439_1_29 | 13/12/2023 | 13/12/2023 | 15381 | ST10 Cplx | A | 6 | 2452 | 4 | 8 | 8 | 8 | 2 |
| 41439_1_65 | 13/12/2023 | 13/12/2023 | 15381 | ST10 Cplx | A | 6 | 2452 | 4 | 8 | 8 | 8 | 2 |
| 41439_1_116 | 13/12/2023 | 13/12/2023 | 15381 | ST10 Cplx | A | 6 | 2452 | 4 | 8 | 8 | 8 | 2 |
| 41439_1_164 | 13/12/2023 | 13/12/2023 | 15381 | ST10 Cplx | A | 6 | 2452 | 4 | 8 | 8 | 8 | 2 |
| 41439_1_113 | 13/12/2023 | 13/12/2023 | 15384 | ST10 Cplx | A | 10 | 11 | 4 | 8 | 1511 | 8 | 2 |
| 42153_1_62 | 13/12/2023 | 13/12/2023 | 15386 | ST10 Cplx | A | 10 | 11 | 4 | 8 | 10 | 13 | 2 |
| 42782_1_334 | 08/01/2024 | 08/01/2024 | 15466 | ST10 Cplx | A | 30 | 11 | 5 | 8 | 7 | 1200 | 2 |
| 42782_2_57 | 08/01/2024 | 08/01/2024 | 15467 | ST10 Cplx | A | 1294 | 11 | 4 | 8 | 8 | 13 | 2 |
| 42782_4_298 | 10/01/2024 | 10/01/2024 | 15475 | ST10 Cplx | A | 6 | 11 | 4 | 8 | 8 | 8 | 1221 |
| 43416_1_222 | 11/01/2024 | 11/01/2024 | 15486 | ST10 Cplx | A | 10 | 11 | 4 | 1954 | 8 | 8 | 2 |
| 42153_2_240 | 14/12/2023 | 14/12/2023 | 11380 | ST131 Cplx | B2 | 53 | 40 | 47 | 1391 | 36 | 28 | 29 |
| 42153_2_298 | 14/12/2023 | 14/12/2023 | 11380 | ST131 Cplx | B2 | 53 | 40 | 47 | 1391 | 36 | 28 | 29 |
| 42782_4_354 | 10/01/2024 | 10/01/2024 | 11380 | ST131 Cplx | B2 | 53 | 40 | 47 | 1391 | 36 | 28 | 29 |
| 41330_8_52 | 13/12/2023 | 13/12/2023 | 15379 | ST131 Cplx | B2 | 53 | 40 | 47 | 13 | 1509 | 28 | 29 |
| 41439_1_248 | 13/12/2023 | 13/12/2023 | 15379 | ST131 Cplx | B2 | 53 | 40 | 47 | 13 | 1509 | 28 | 29 |
| 42756_3_213 | 15/12/2023 | 15/12/2023 | 15379 | ST131 Cplx | B2 | 53 | 40 | 47 | 13 | 1509 | 28 | 29 |
| 43416_1_30 | 11/01/2024 | 11/01/2024 | 15379 | ST131 Cplx | B2 | 53 | 40 | 47 | 13 | 1509 | 28 | 29 |
| 42782_2_276 | 10/01/2024 | 10/01/2024 | 7322 | ST155 Cplx | | 6 | 1007 | 14 | 16 | 24 | 8 | 14 |
| 42782_2_42 | 08/01/2024 | 08/01/2024 | 8679 | ST155 Cplx | | 6 | 4 | 14 | 16 | 748 | 8 | 14 |
| 41439_1_255 | 13/12/2023 | 13/12/2023 | 8744 | ST155 Cplx | | 6 | 4 | 14 | 989 | 24 | 8 | 14 |
| 42756_3_19 | 14/12/2023 | 14/12/2023 | 8744 | ST155 Cplx | | 6 | 4 | 14 | 989 | 24 | 8 | 14 |
| 42756_3_57 | 14/12/2023 | 14/12/2023 | 8744 | ST155 Cplx | | 6 | 4 | 14 | 989 | 24 | 8 | 14 |
| 42756_3_148 | 15/12/2023 | 15/12/2023 | 8744 | ST155 Cplx | | 6 | 4 | 14 | 989 | 24 | 8 | 14 |
| 42153_1_78 | 14/12/2023 | 14/12/2023 | 11189 | ST155 Cplx | | 6 | 1181 | 14 | 16 | 24 | 8 | 14 |
| 42153_1_245 | 14/12/2023 | 14/12/2023 | 11189 | ST155 Cplx | | 6 | 1181 | 14 | 16 | 24 | 8 | 14 |
| 42153_1_275 | 14/12/2023 | 14/12/2023 | 11189 | ST155 Cplx | | 6 | 1181 | 14 | 16 | 24 | 8 | 14 |
| 42153_2_341 | 14/12/2023 | 14/12/2023 | 11189 | ST155 Cplx | | 6 | 1181 | 14 | 16 | 24 | 8 | 14 |
| 42601_1_320 | 14/12/2023 | 14/12/2023 | 11189 | ST155 Cplx | | 6 | 1181 | 14 | 16 | 24 | 8 | 14 |
| 42782_1_32 | 08/01/2024 | 08/01/2024 | 11189 | ST155 Cplx | | 6 | 1181 | 14 | 16 | 24 | 8 | 14 |
| 42782_1_210 | 08/01/2024 | 08/01/2024 | 11189 | ST155 Cplx | | 6 | 1181 | 14 | 16 | 24 | 8 | 14 |
| 42782_2_101 | 08/01/2024 | 08/01/2024 | 11189 | ST155 Cplx | | 6 | 1181 | 14 | 16 | 24 | 8 | 14 |
| 42782_2_109 | 08/01/2024 | 08/01/2024 | 11189 | ST155 Cplx | | 6 | 1181 | 14 | 16 | 24 | 8 | 14 |
| 42782_2_116 | 08/01/2024 | 08/01/2024 | 11189 | ST155 Cplx | | 6 | 1181 | 14 | 16 | 24 | 8 | 14 |
| 42782_2_119 | 08/01/2024 | 08/01/2024 | 11189 | ST155 Cplx | | 6 | 1181 | 14 | 16 | 24 | 8 | 14 |
| 42782_2_215 | 08/01/2024 | 08/01/2024 | 11189 | ST155 Cplx | | 6 | 1181 | 14 | 16 | 24 | 8 | 14 |
| 42782_4_217 | 10/01/2024 | 10/01/2024 | 15472 | ST155 Cplx | | 6 | 2465 | 14 | 16 | 24 | 8 | 14 |
| 41439_1_36 | 13/12/2023 | 13/12/2023 | 15383 | ST156 Cplx | AxB1 | 1861 | 29 | 32 | 16 | 11 | 8 | 44 |
| 42153_2_294 | 14/12/2023 | 14/12/2023 | 9816 | ST206 Cplx | AxB1 | 6 | 7 | 856 | 1 | 8 | 18 | 2 |
| 41439_1_104 | 13/12/2023 | 13/12/2023 | 10822 | ST206 Cplx | AxB1 | 1110 | 7 | 5 | 1 | 8 | 18 | 2 |
| 42756_3_1 | 14/12/2023 | 14/12/2023 | 10822 | ST206 Cplx | AxB1 | 1110 | 7 | 5 | 1 | 8 | 18 | 2 |
| 42756_4_183 | 15/12/2023 | 15/12/2023 | 9347 | ST23 Cplx | | 6 | 4 | 14 | 1084 | 20 | 62 | 7 |
| 43416_1_192 | 11/01/2024 | 11/01/2024 | 15487 | ST23 Cplx | B1 | 6 | 4 | 33 | 132 | 20 | 12 | 949 |
| 42782_1_322 | 08/01/2024 | 08/01/2024 | 3385 | ST38 Cplx | D | 4 | -43 | 2 | 25 | 5 | 5 | 19 |
| 42601_1_242 | 14/12/2023 | 14/12/2023 | 15389 | ST38 Cplx | D | 271 | 26 | 39 | 25 | 1513 | 31 | 19 |
| 42756_4_115 | 15/12/2023 | 15/12/2023 | 15401 | ST394 Cplx | D | 21 | 35 | 1539 | 52 | 5 | 5 | 4 |
| 43416_1_102 | 11/01/2024 | 11/01/2024 | 15401 | ST394 Cplx | D | 21 | 35 | 1539 | 52 | 5 | 5 | 4 |
| 41439_1_250 | 13/12/2023 | 13/12/2023 | 12637 | ST469 Cplx | AxB1 | 109 | 65 | 244 | 1 | 9 | 13 | 14 |
| 41330_8_224 | 13/12/2023 | 13/12/2023 | 8881 | ST648 Cplx | | 92 | 4 | 87 | 96 | 70 | 13 | 2 |
| 41439_1_213 | 13/12/2023 | 13/12/2023 | 8881 | ST648 Cplx | | 92 | 4 | 87 | 96 | 70 | 13 | 2 |
| 42476_3_9 | 14/12/2023 | 14/12/2023 | 8881 | ST648 Cplx | | 92 | 4 | 87 | 96 | 70 | 13 | 2 |
| 42756_3_261 | 15/12/2023 | 15/12/2023 | 8881 | ST648 Cplx | | 92 | 4 | 87 | 96 | 70 | 13 | 2 |
| 43416_1_173 | 11/01/2024 | 11/01/2024 | 8881 | ST648 Cplx | | 92 | 4 | 87 | 96 | 70 | 13 | 2 |
| 43416_1_238 | 11/01/2024 | 11/01/2024 | 8881 | ST648 Cplx | | 92 | 4 | 87 | 96 | 70 | 13 | 2 |
| 42756_3_40 | 14/12/2023 | 14/12/2023 | 7937 | | | 6 | 4 | 15 | 16 | 11 | 8 | 6 |
| 41439_1_154 | 13/12/2023 | 13/12/2023 | 8131 | | D | 5 | 3 | 2 | 6 | 45 | 5 | 4 |
| 43416_1_41 | 11/01/2024 | 11/01/2024 | 8131 | | D | 5 | 3 | 2 | 6 | 45 | 5 | 4 |
| 42756_3_36 | 14/12/2023 | 14/12/2023 | 8330 | | | 806 | 1096 | 701 | 520 | 401 | 40 | 350 |
| 42756_3_153 | 15/12/2023 | 15/12/2023 | 8577 | | A | 8 | 7 | 728 | 220 | 8 | 8 | 2 |
| 43416_1_78 | 11/01/2024 | 11/01/2024 | 8577 | | A | 8 | 7 | 728 | 220 | 8 | 8 | 2 |
| 41330_8_277 | 13/12/2023 | 13/12/2023 | 9439 | | A | 10 | 11 | 1 | 8 | 12 | 18 | 2 |
| 41430_160 | 13/12/2023 | 13/12/2023 | 9523 | | ABD | 12 | 371 | 176 | 12 | 1 | 2 | 2 |
| 41439_1_267 | 13/12/2023 | 13/12/2023 | 9523 | | ABD | 12 | 371 | 176 | 12 | 1 | 2 | 2 |
| 43416_1_20 | 11/01/2024 | 11/01/2024 | 9523 | | ABD | 12 | 371 | 176 | 12 | 1 | 2 | 2 |
| 41330_8_185 | 13/12/2023 | 13/12/2023 | 10391 | | | 218 | 371 | 53 | 140 | 247 | 2 | 216 |
| 43416_1_235 | 11/01/2024 | 11/01/2024 | 12137 | | | 6 | 4 | 1159 | 102 | 9 | 73 | 682 |
| 42782_3_163 | 10/01/2024 | 10/01/2024 | 12290 | | ABD | 826 | 186 | 54 | 10 | 1 | 35 | 47 |
| 41439_1_11 | 13/12/2023 | 13/12/2023 | 12400 | | A | 8 | 1873 | 1 | 8 | 8 | 18 | 6 |
| 42153_1_254 | 14/12/2023 | 14/12/2023 | 12569 | | D | 200 | 3 | 174 | 6 | 1187 | 5 | 191 |
| 42601_1_214 | 14/12/2023 | 14/12/2023 | 12569 | | D | 200 | 3 | 174 | 6 | 1187 | 5 | 191 |
| 42782_1_67 | 08/01/2024 | 08/01/2024 | 12569 | | D | 200 | 3 | 174 | 6 | 1187 | 5 | 191 |
| 42782_1_115 | 08/01/2024 | 08/01/2024 | 12569 | | D | 200 | 3 | 174 | 6 | 1187 | 5 | 191 |
| 42782_1_179 | 08/01/2024 | 08/01/2024 | 12569 | | D | 200 | 3 | 174 | 6 | 1187 | 5 | 191 |
| 42782_3_52 | 09/01/2024 | 09/01/2024 | 12569 | | D | 200 | 3 | 174 | 6 | 1187 | 5 | 191 |
| 42782_3_72 | 09/01/2024 | 09/01/2024 | 12569 | | D | 200 | 3 | 174 | 6 | 1187 | 5 | 191 |
| 42782_3_104 | 10/01/2024 | 10/01/2024 | 12569 | | D | 200 | 3 | 174 | 6 | 1187 | 5 | 191 |
| 42782_3_211 | 10/01/2024 | 10/01/2024 | 12569 | | D | 200 | 3 | 174 | 6 | 1187 | 5 | 191 |
| 42782_3_303 | 10/01/2024 | 10/01/2024 | 12569 | | D | 200 | 3 | 174 | 6 | 1187 | 5 | 191 |
